# Visualization of *trans* homophilic interaction of clustered protocadherin in neurons

**DOI:** 10.1101/2023.04.14.536980

**Authors:** Natsumi Hoshino, Takashi Kanadome, Mizuho Itoh, Ryosuke Kaneko, Yukiko U. Inoue, Takayoshi Inoue, Takahiro Hirabayashi, Masahiko Watanabe, Tomoki Matsuda, Takeharu Nagai, Etsuko Tarusawa, Takeshi Yagi

**Author notes:** Corresponding author: Takeshi Yagi, **Email:**. **Author Contributions:** Conceptualization, N.H. and T.Y.; Formal analysis, N.H. and K.T.; Investigation, N.H., T.K., M.U., T. H., Y.U.I., T.I., and E.T.; Resources, R.K., T.H., and M.W.; Writing – Original Draft, N.H., E.T., and T. Y.; Supervision, T.M., T.N., and T.Y.; Funding Acquisition, T.K., E.T., and T.Y. **Competing Interest Statement:** The authors declare no competing interests.

## Abstract

Clustered protocadherin (Pcdh) functions as a cell recognition molecule through the homophilic interaction in CNS. However, its interactions have yet not been visualized in neurons. We previously reported PcdhγB2-FRET probes to be applicable only for cell lines. Herein, we newly designed PcdhγB2-FRET probes by fusing FRET donor and acceptor fluorescent proteins to a single PcdhγB2 molecule and succeeded in visualizing PcdhγB2 homophilic interaction in cultured hippocampal neurons. The γB2-FRET probe localized in the soma and neurites, and FRET signals were observed at contact sites between neurites and eliminated by EGTA addition. Live imaging revealed that the FRET-negative γB2 signals were rapidly moving along neurites and soma, whereas the FRET-positive signals remained in place. We observed that the γB2 proteins at synapses rarely interact homophilically. The γB2-FRET probe would allow us to elucidate the function of the homophilic interaction and the cell recognition mechanism.

**Significance Statement:** We visualize the Pcdh homophilic interaction using a novel FRET-based probe, and reveal that the homophilically interacting Pcdh proteins are found at contact sites between the neurites and roots of neurites from the soma, and are stable at a location. Additionally, in neurons, Pcdh proteins are located at synapses but rarely interact homophilically.

## Introduction

Neural networks are formed in billions of neurons. Cell surface proteins are required for recognizing surrounding neurites, including self- and non-self-recognition (1). Clustered protocadherin (Pcdh) proteins, which comprise 58 isoforms in mice, are thought to provide neuron identity through their stochastic expression and homophilic interaction (2–5). Stochastic expression in neurons has been revealed by single-cell RT-PCR (6–8) and single-cell RNA sequencing in cortical and olfactory sensory neurons (9–11).

The Pcdh homophilic interaction has been studied using cell aggregation assay (3, 4, 12), structural information of *trans* dimers (13–15), and biophysical measurements (3, 14, 16). In neurons, homophilic interactions are implicated through self-crossing phenotypes in Pcdhγ KO mice (17–19) and overexpressing matched-/mismatched-isoforms (20, 21); however, where and when the Pcdh homophilic interactions occur in neurons remains unknown.

We previously reported a FRET probe for visualizing Pcdh homophilic interactions in cultured cell lines (22). Previous studies have shown that Pcdhγ proteins are localized at the contact sites of neurites and synapses (23–26).

In the present study, we aimed to develop a novel FRET probe in which CFP and YFP were fused to a single Pcdh molecule. We successfully visualized the Ca2+-dependent Pcdh homophilic interaction in neurons and generated Pcdhγ-floxed mice and PcdhγB2 FRET mice to determine whether the FRET probe functions as a Pcdhγ protein in neurons *in vivo*. To the best of our knowledge, we provide the first evidence that Pcdhγ proteins show homophilic interactions in neurons at the contact sites of neurites and that Pcdhγ proteins at synapses rarely interact homophilically.

## Results

### γB2ΔICD-EC1-CFP-EC5-YFP visualizes the γB2 homophilic interaction as FRET

Previously, we produced a set of FRET probes comprising γB2-EC1-CFP (CFP fused to extracellular domain 1 of PcdhγB2) and γB2-EC5-YFP (YFP fused to extracellular domain 5 of PcdhγB2): separated γB2-FRET probes (Figure 1A) (22). We prepared γB2-FRET probes with the intracellular domains (ICD) deleted because ICD deletion has been reported to result in efficient localization to the plasma membrane (23). To observe FRET signals, we performed ratio imaging. Wherein CFP was excited with a 405 nm laser and the fluorescence of the CFP and YFP emission wavelengths were collected simultaneously. The former and latter are known as the CFP and FRET channels, respectively. The YFP channel, which shows the fluorescence of YFP using a YFP excitation laser (514 nm laser), was used to observe the localization of YFP. FRET was visualized as an increase in the FRET/CFP ratio.

**Figure 1.**
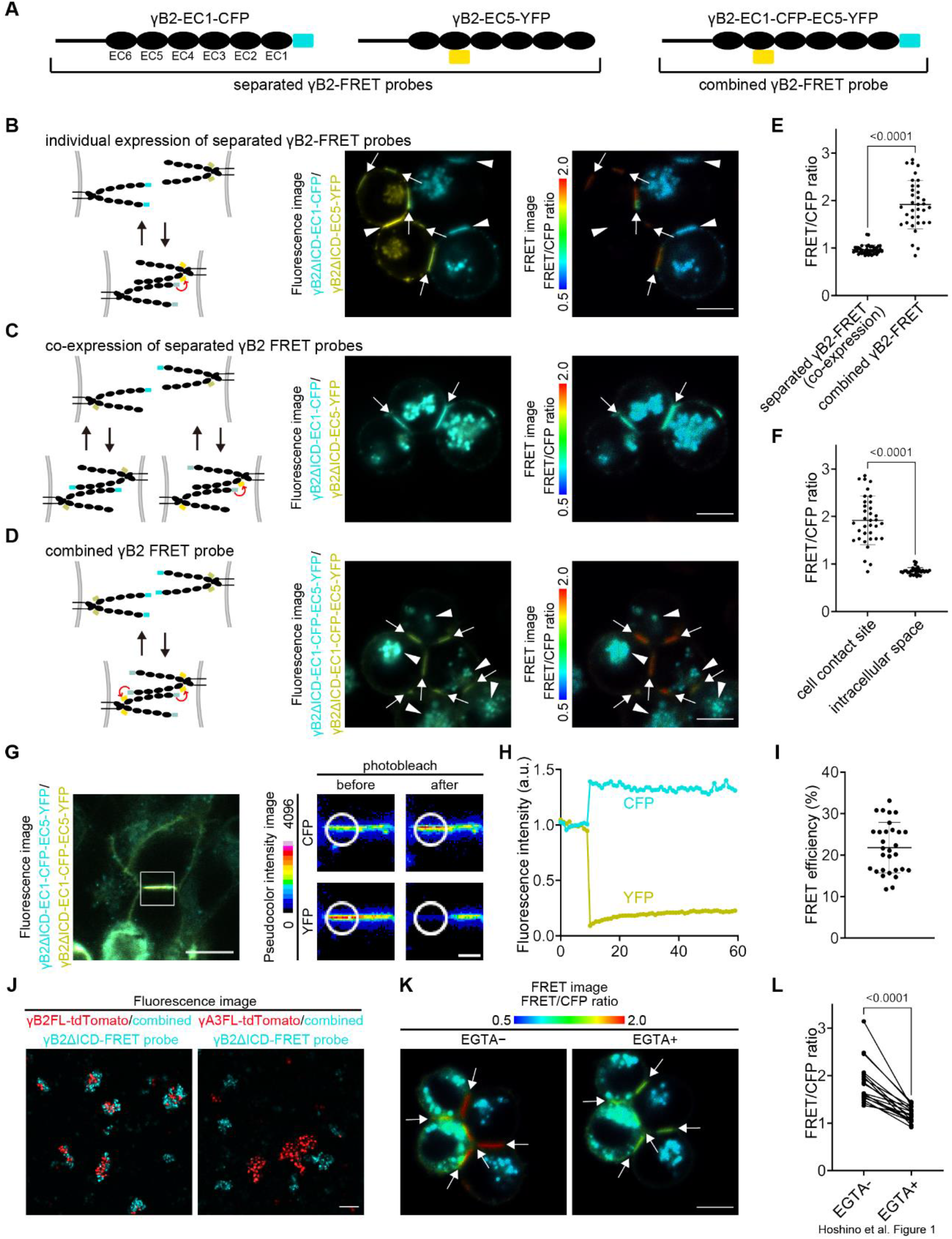
Combined γB2-FRET probe visualizes homophilic interaction. (A) Schematics of the γB2-FRET probes used in this study. Black ovals represent the extracellular domains of PcdhγB2. Cyan and yellow ovals represent CFP and YFP, respectively. (B) Schematics of the non-interacting separated γB2-FRET probes and interacting separated γB2-FRET probes when γB2-EC1-CFP and γB2-EC5-YFP are individually expressed in cells (left). Fluorescence images showing the localization of γB2ΔICD-EC1-CFP and γB2ΔICD-EC1-YFP in K562 cells (middle). A ratio image showing FRET as an increased FRET/CFP ratio (right). Arrows indicate the contact sites between a γB2ΔICD-EC1-CFP expressing cell and a γB2ΔICD-EC5-YFP expressing cell. Arrowheads indicate the contact sites between two γB2ΔICD-EC1-CFP expressing cells or two γB2ΔICD-EC5-YFP expressing cells. Scale bar, 10 µm. (C) Schematics of the non-interacting separated γB2-FRET probes and interacting separated γB2-FRET probes when γB2-EC1-CFP and γB2-EC5-YFP are coexpressed in each cell (left). Fluorescence images showing the localization of γB2ΔICD-EC1-CFP and γB2ΔICD-EC1-YFP in K562 cells (middle). A ratio image showing FRET as an increased FRET/CFP ratio (right). Arrows indicate the contact sites between two γB2ΔICD-EC1-CFP and γB2ΔICD-EC5-YFP expressing cells. Scale bar, 10 µm. (D) Schematics of the non-interacting combined γB2-FRET probes and interacting combined γB2-FRET probes (left). Fluorescence images showing the localization of γB2ΔICD-EC1-CFP-EC5-YFP in K562 cells (middle). A ratio image showing FRET as an increased FRET/CFP ratio (right). Arrows indicate the contact sites between two γB2ΔICD-EC1-CFP-EC5-YFP expressing cells. Arrowheads indicate the intracellular spaces of γB2ΔICD-EC1-CFP-EC5-YFP expressing cells. Scale bar, 10 µm. (E) Quantification of the FRET/CFP ratio at cell contact sites of co-expression of separated γB2ΔICD-FRET probe and the cell contact sites of combined γB2ΔICD-FRET probe (n = 5; mean ± SD; Welch’s t-test). (F) Quantification of the FRET/CFP ratio at cell contact sites and intracellular spaces of combined γB2ΔICD-FRET probe (n = 5; mean ± SD; Welch’s t-test). The data for cell contact sites are the same as in Figure 1E. (G) HEK293T cells expressing combined γB2ΔICD-FRET probe. The right panels show magnified images of the white squared region in the left panel. The region of the white circle was acceptor photobleached and analyzed the fluorescence intensity for Figure 1H. Scale bars, 10 µm (left) and 2 µm (right). (H) Quantification of CFP channel and YFP channel fluorescence intensity before and after acceptor photobleaching. (I) Quantification of FRET efficiency of the combined γB2ΔICD-FRET probe using acceptor photobleaching (n = 30; mean ± SD). (J) K562 cell aggregation using the matched and mismatched isoforms for the combined γB2ΔICD-FRET probe. Scale bar, 100 µm. (K) A ratio image of combined γB2ΔICD-FRET probe-expressing K562 cells before and after EGTA addition. Arrows indicate cell contact sites. Scale bar, 10 µm. (L) Quantification of FRET/CFP ratio changes via EGTA addition (n = 5; Paired t-test).

The separated FRET probes were individually transfected with γB2ΔICD-EC1-CFP and γB2ΔICD-EC5-YFP into each cell line (Figure 1B, arrows). However, this strategy cannot visualize all γB2 homophilic interactions because CFP-CFP and YFP-YFP proximities were not visualized with FRET (Figure 1B, arrowheads). To avoid this problem, we transfected γB2ΔICD-EC1-CFP and γB2ΔICD-EC5-YFP into each cell line. However, as reported in our previous study, the cell contact sites of the cotransfected cells did not show any FRET signals despite the γB2 homophilic interaction (22) (Figure 1C).

Based on the γB2 *trans* dimer crystal structure (14), CFP and YFP can be fused to a single γB2 molecule, therefore we generated a γB2-EC1-CFP-EC5-YFP: combined γB2-FRET probe (Figure 1A). Ratio imaging showed that the combined γB2ΔICD-FRET probe could visualize the γB2 homophilic interactions (Figure 1D). The combined γB2ΔICD-FRET probes in intracellular spaces did not show FRET signals (Figures 1D and 1E). These results reveal that the combined γB2ΔICD-FRET probes can be used to visualize the *trans* homophilic interaction of γB2. Acceptor photobleaching was used to quantify the FRET efficiency. In this experiment, YFP was photobleached and the fluorescence recovery of CFP was verified (Figures 1G and 1H). The combined γB2ΔICD-FRET probe in HEK293T cells revealed a FRET efficiency of 21.8% ± 6.06% (Figure 1I, mean ± SD). This is slightly lower than the separated γB2ΔICD-FRET probes (27.4 ± 5.67%) (22), but is high enough to analyze the homophilic interaction of γB2.

It is possible that the increased FRET/CFP ratio at the cell contact sites is due to the change in environmental factors between the extracellular and intracellular regions, that a *cis* interaction at the cell contact sites could show FRET signals, and that FRET occurs intramolecularly at the cell contact sites. To exclude these possibilities, we co-cultured combined γB2ΔICD-FRET probe-expressing K562 cells with γB2ΔICD-mCherry-expressing K562 cells (Figures S1A and S1B).

Ratio imaging revealed that FRET signals were observed only at the cell contact sites between the two combined γB2ΔICD-FRET probe-expressing cells, whereas no FRET signal was observed between the combined γB2ΔICD-FRET probe-expressing cell and the γB2ΔICD-mCherry-expressing cell (Figures S1B and S1C). These results showed that the FRET signals with the combined γB2ΔICD-FRET probe successfully reflected the homophilic interaction of γB2. To confirm the specificity of the homophilic interaction of the combined γB2ΔICD-FRET probe, we performed K562 cell aggregation assays. The cells expressing the combined γB2ΔICD-FRET probe aggregated with the cells expressing γB2FL (full-length)-tdTomato, whereas they segregated from those expressing γA3FL-tdTomato (Figure 1J). These results indicate that the combined γB2ΔICD-FRET probe maintains homophilic binding specificity.

To determine whether the CFP and YFP of the extracellular domains of γB2 interrupt the *cis* interaction with other Pcdh isoforms, we observed the ability to deliver Pcdhα4, which requires *cis* interaction with Pcdhβ or Pcdhγ isoforms to express on the cell membrane. HEK293T cells were transfected with α4ΔICD-mCherry alone, cotransfected with γB2ΔICD-Venus, or a combined γB2ΔICD-FRET probe. In γB2ΔICD-Venus, Venus is fused to the C-terminus of γB2ΔICD. When the γB2ΔICD-Venus or combined γB2ΔICD-FRET probe was co-expressed, α4ΔICD-mCherry proteins were localized to the plasma membrane, whereas they localized at intracellular spaces when α4ΔICD-mCherry proteins were expressed alone (Figure S1D). Co-immunoprecipitation using K562 cells also showed that the combined γB2FL-FRET probe interacted in *cis* with α4 and γB2FL-Venus, whose Venus was fused to the C-terminal of γB2FL (Figure S1E). These results suggest that the combined γB2ΔICD-FRET and the combined γB2FL-FRET probe maintain their *cis* interaction properties with α4.

As reported for the separated γB2-FRET probes (22), the FRET signals at the cell contact sites of the combined γB2ΔICD-FRET probe-expressing cells decreased upon EGTA addition (Figures 1K and 1L). These results indicate that the homophilic interaction of the γB2-FRET probe is extracellular calcium-dependent.

### Direct excitation of the acceptor is normalized in a combined γB2-FRET probe

Phototoxicity is a matter of concern when FRET signals are observed in neurons. Therefore, for observation in neurons, we planned to use a 458 nm laser instead of the 405 nm laser used in Figure 1. With ratio imaging, the YFP-rich region compared with CFP can be visualized as if FRET occurred because the 458 nm laser can directly excite YFP. To verify the pseudo-positive FRET signals caused by the direct excitation of YFP, we prepared negative controls for the separated and combined γB2-FRET probe, whose YFP was fused in the C-terminal region of γB2ΔICD. Therefore, the cell contact sites in the negative control should not cause FRET. With the separated C-terminal probe, the FRET/CFP ratio at cell contact sites was higher than that in the intracellular spaces of CFP-expressing cells because of the direct excitation and unequal molar ratio of CFP and YFP (Figures 2A, 2C (i)–(iv), and 2D). This would be a problem in neurons since the contact sites and intracellular spaces of thin neurites cannot be identified, unlike the obvious contact sites and intracellular spaces of K562 cells, and since the ratio of the molecular numbers of γB2-EC1-CFP and γB2-EC5-YFP from two neurites in each pixel varies and such inequality is difficult to normalize accurately.

**Figure 2.**
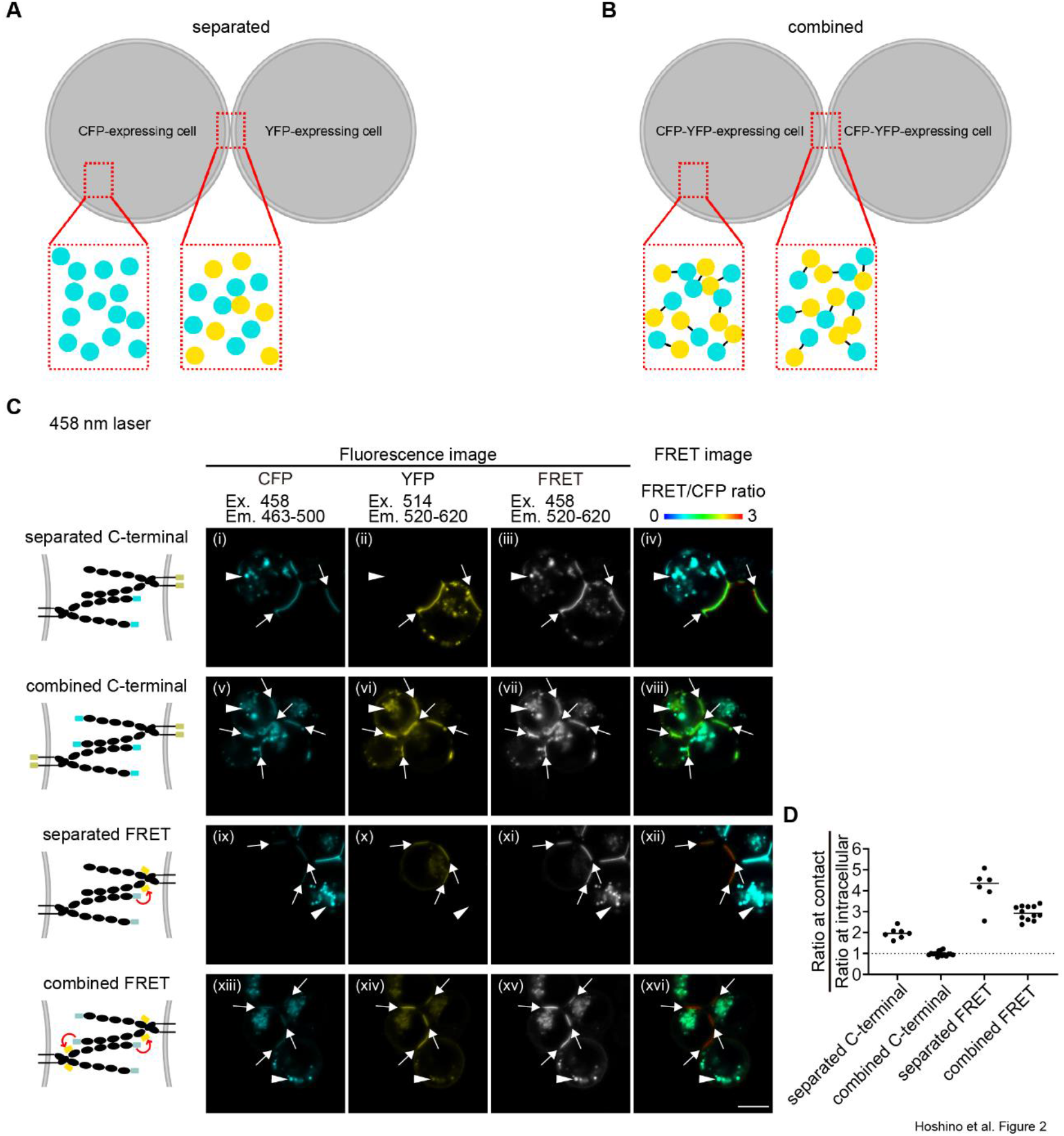
Combined γB2-FRET probe visualizes the homophilic interaction using a 458 nm laser. (A) Schematic of uneven distribution of CFP and YFP using the separated γB2-FRET probes. If the laser for CFP excitation also excites YFP, FRET/CFP ratio will increase at cell contact sites when compared with the one at intracellular spaces in the CFP-expressing cells even when there is no FRET signal at the cell contact sites. (B) Schematic of even distribution of CFP and YFP using the combined γB2-FRET probes. Even if the laser excites YFP, FRET/CFP ratio is the same between the cell contact sites and intracellular spaces. (C) Ratio images of the separated and combined γB2ΔICD-C-terminal negative control and γB2ΔICD-FRET probes expressed in K562 cells with a 458 nm laser. Abbreviations: Em, emission; Ex, excitation. Arrows indicate the cell contact sites. Arrowheads indicate intracellular spaces. Scale bar, 10 µm. (D) Quantification of the FRET/CFP ratio at cell contact sites normalized with the one at intracellular spaces (n = 3; mean).

However, the combined C-terminal probe did not show pseudo-positive FRET signals because CFP and YFP were equimolar, and the ratio of direct excitation of YFP was constant to CFP fluorescence (Figures 2B, 2C (v)–(viii) and 2D). In the separated and combined γB2ΔICD-FRET probes, they successfully showed that higher ratios were observed at the cell contact sites than in the intracellular spaces (Figure 2C (ix)–(xvi) and 2D). If phototoxicity is not considered, the 405 nm laser can be used for both the separated and γB2-FRET probes because almost no YFP is directly excited by a 405 nm laser (Figures S2A and S2B). These results suggest that the combined γB2-FRET probe can be used with a 458 nm laser.

### FRET+ combined γB2-FRET probe in cultured hippocampal neurons is stable at sites

To visualize the γB2 homophilic interaction in neurons, we prepared cultured hippocampal neurons from mice at E18.5 and transfected combined γB2FL-FRET probes. We discriminated autofluorescence using spectral imaging, followed by linear mixing since we found that in rare neurons, non-transfected neurons showed dotted fluorescence. Linear unmixing is a mathematical technique used to decompose the spectra of mixed samples into pure components. We obtained three reference spectra: autofluorescence from a cultured hippocampal neuron, FRET− from the intracellular space of K562 cells expressing the combined γB2ΔICD-FRET probe, and FRET+ from the cell contact sites of K562 cells expressing the combined γB2ΔICD-FRET probe (Figure S3A). Using K562 cells, the linear unmixed image showed that FRET+ signals were found at the cell contact sites, and FRET− signals were found in the intracellular spaces (Figure S3B). EGTA addition decreased the FRET+ ratio at cell contact sites (Figures S3B and S3C). These results suggest that spectral imaging followed by linear unmixing is applicable for analyzing FRET signals.

To apply linear unmixing in cultured hippocampal neurons, samples were excited with a 458 nm laser, and the fluorescence was collected using spectrum imaging. The fluorescence was linearly unmixed with autofluorescence, FRET−, and FRET+ (Figure S3B) (See STAR Methods). Pcdhγ proteins show a dot-like localization (2, 23–25, 27). We transfected the plasmid encoding the combined γB2FL-FRET probe into cultured hippocampal neurons along with mCherry or iRFP670 to visualize cell morphology (Figure 3A). As expected, the combined γB2FL-FRET probes showed dot-like localization. FRET− signals were found in the soma and neurites, whereas FRET+ signals were detected at the contact sites between neurites and the roots of neurites from the soma (Figure 3A). Live imaging revealed that while the γB2 proteins showing FRET+ signals and most showing FRET− signals were immobile, some γB2 proteins showing FRET− signals were moving (Figures 3B and 3C). This indicates that the homophilically interacting γB2 proteins are stable at the contact sites, and γB2 proteins that undergo trafficking are not homophilically interacting.

**Figure 3.**
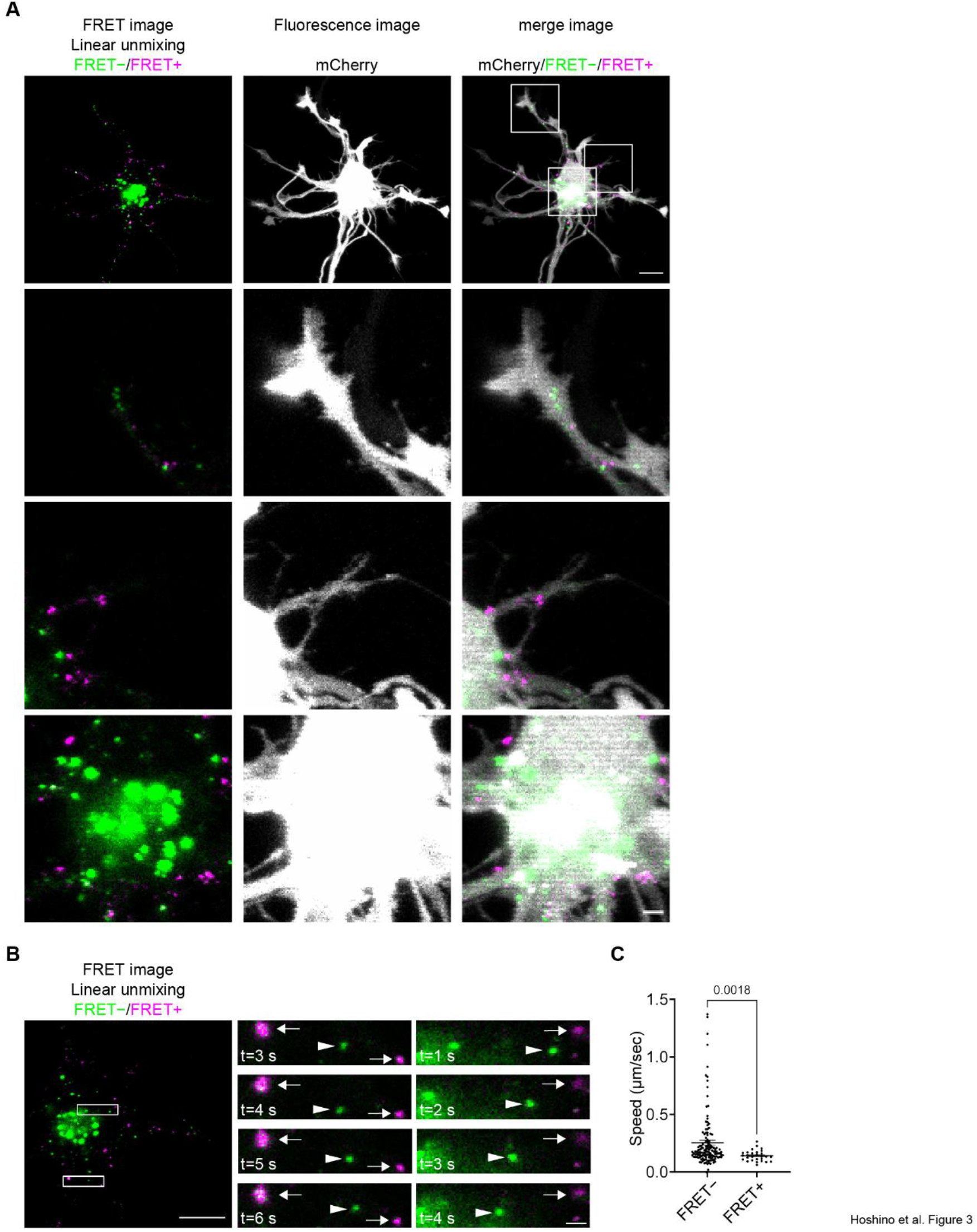
γB2-FRET signals are observed in neurons. (A) Linear unmixed images of a cultured hippocampal neuron. mCherry was transfected to visualize the cell morphology. The bottom 3 fields of view are magnified images in white rectangles in the top panels. Scale bars, 10 µm for the top, and 2 µm for the bottom. (B) Time-lapse imaging of the combined γB2FL-FRET probes. Arrows indicate combined γB2FL-FRET probes showing FRET+ signals. Arrowheads indicate γB2-FRET probes showing FRET− signals. Scale bar, 10 µm for the left panel, 2 µm for the right panel. (C) Quantification of the speed of each dot (n = 6; mean ± SEM; Mann Whitney test).

### Conditionally overexpressed combined γB2FL-FRET probe rescues Pcdhγ conditional knockout phenotype

For further analysis in neurons, we investigated whether the combined γB2FL-FRET probe functions as Pcdhγ in neurons. Little is known about the function of a single Pcdhγ isoform. Pcdhγ deletion disrupts the self-avoidance of dendrites, and this phenotype can be rescued by the overexpression of a single Pcdhγ isoform (19). To evaluate the function of the combined γB2FL-FRET probe as a single Pcdhγ isoform, we investigated whether this self-crossing phenotype could be rescued by the combined γB2FL-FRET probe in Purkinje cells. Accordingly, we generated Pcdhγ-floxed mice and γB2-FRET mice, which conditionally knocked out Pcdhγ and overexpressed the combined γB2FL-FRET probe in a Cre-dependent manner, respectively (Figure S4).

Three genotypes were established: control mice (*PV*^*PV-Cre/WT*^: *Pcdhγ*^*flox/WT*^; *γB2-FRET*^*−/−*^ or PV^WT/WT^; *Pcdhγ*^*flox/flox*^; γB2-FRET^−/−^), Pcdhγ cKO + γB2-FRET cOE mice (*PV*^*PV-Cre/WT*^; *Pcdhγ*^*flox/flox*^; *γB2-FRET*^*Tg/Tg*^), and Pcdhγ cKO mice (*PV*^*PV-Cre/WT*^; *Pcdhγ*^*flox/flox*^; *γB2-FRET*^*−/−*^). The neurites of a single Purkinje cell were visualized and the number of self-crossings was counted. In ΔPcdhγ mice, the dendrites were multiplanar rather than monoplanar in control mice, and more self-crossings were observed than in control mice. In Pcdhγ cKO + γB2-FRET cOE mice, the dendrites were monoplanar, as observed in control mice, and the self-crossing phenotype of Pcdhγ cKO was rescued by the expression of the combined γB2FL-FRET probe (Figures 4A and 4B). These results suggest that a single Pcdhγ isoform can rescue the self-crossing phenotype in Purkinje cells and that the combined γB2FL-FRET probe possesses a self-avoiding function that native PcdhγB2 should possess.

**Figure 4.**
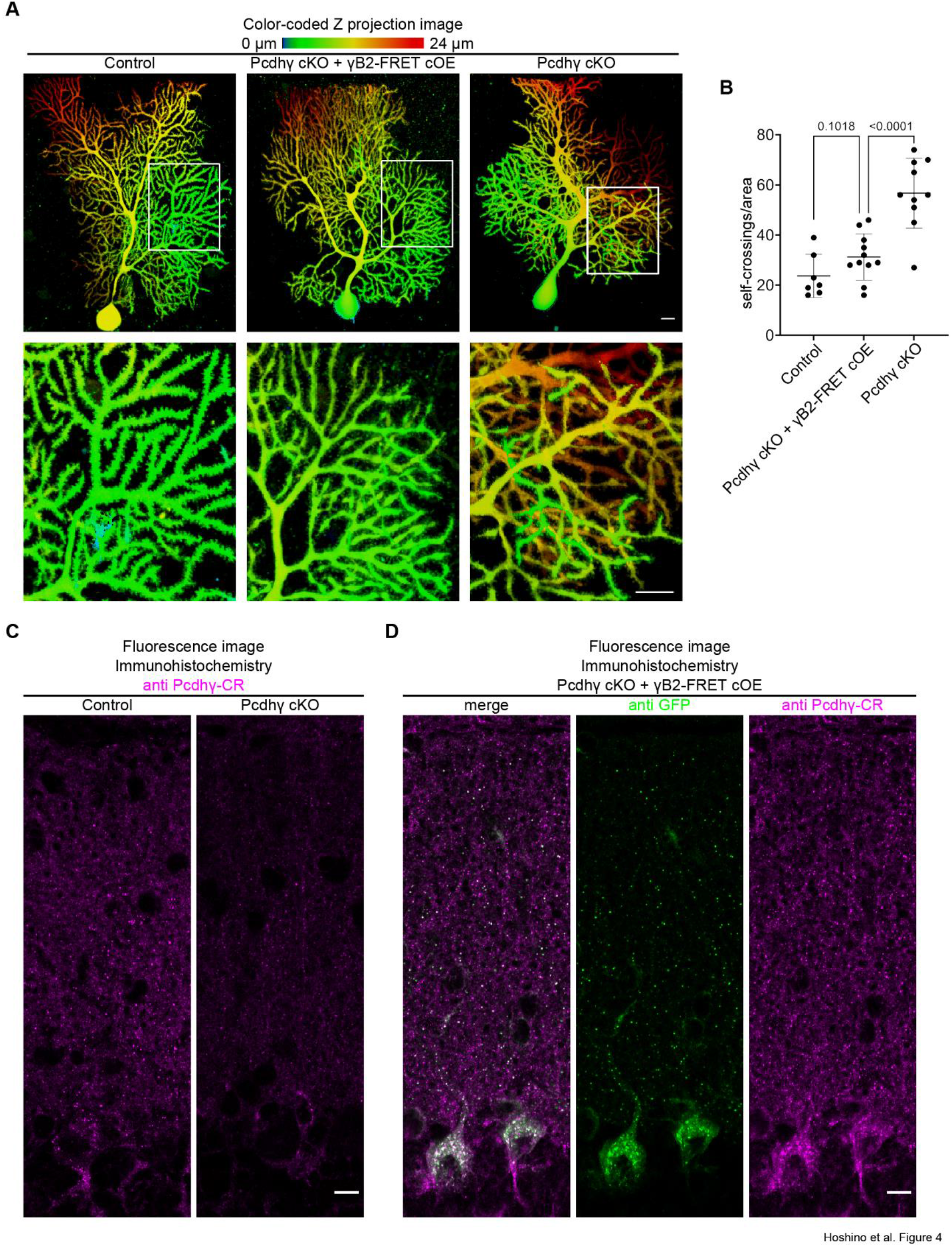
Conditionally overexpressed combined γB2-FRET probe rescues Pcdhγ conditional knockout phenotype. (A) Dendrites of a single Purkinje cell were visualized and color-coded according to the z-axis. The bottom panels are the magnified images in white boxes in the top panels. Scale bars, 10 µm. (B) Quantification showing self-crossings in each genotype (n = 7 for control, n=11 for Pcdhγ cKO + γB2-FRET cOE, and n=10 for Pcdhγ cKO; mean ± SD; Unpaired t-test). (C) Immunohistochemistry with an anti Pcdhγ antibody in the cerebellum. The fluorescence decreased in the Pcdhγ cKO mice when compared to control mice. Scale bar, 10 µm. (D) Immunohistochemistry with an anti-GFP antibody and an anti Pcdhγ antibody in the cerebellum. The anti-GFP signal colocalized with the anti Pcdhγ signal. Scale bar, 10 µm.

Immunohistochemistry using an antibody for the Pcdhγ constant region (Pcdhγ CR) to detect all Pcdhγ isoforms revealed that endogenous Pcdhγ proteins were expressed in the cerebellar molecular layer and the signal decreased in Pcdhγ cKO mice when PV-Cre was expressed (Figure 4C). An antibody against GFP was used to detect CFP and YFP in the combined γB2FL-FRET probe. In Pcdhγ cKO + γB2-FRET cOE mice, overexpression of the combined γB2FL-FRET probe was confirmed, and the combined γB2FL-FRET probe was localized in the cerebellar molecular layer, similar to endogenous Pcdhγ in control mice (Figures 4C and 4D).

### Homophilic interaction of exogenously overexpressed γB2 isoform is interrupted with endogenous Pcdhs

A cell aggregation assay using K562 cells showed that mismatched isoforms inhibited cell aggregation, although matched isoforms were expressed between cells (4). To investigate the spatiotemporal features of the γB2 homophilic interaction and the effects of endogenous Pcdh isoforms, we prepared cultured hippocampal neurons from WT and Δαβγ mice, which lack all clustered Pcdh isoforms (28). We observed γB2-FRET signals during development.

To analyze the γB2-FRET signals, we quantified the fluorescence intensity unmixed for FRET+ and compared it to the total fluorescence intensity from the combined γB2-FRET probe (FRET− + FRET+), as shown in Figure 5A (See STAR Methods).

**Figure 5.**
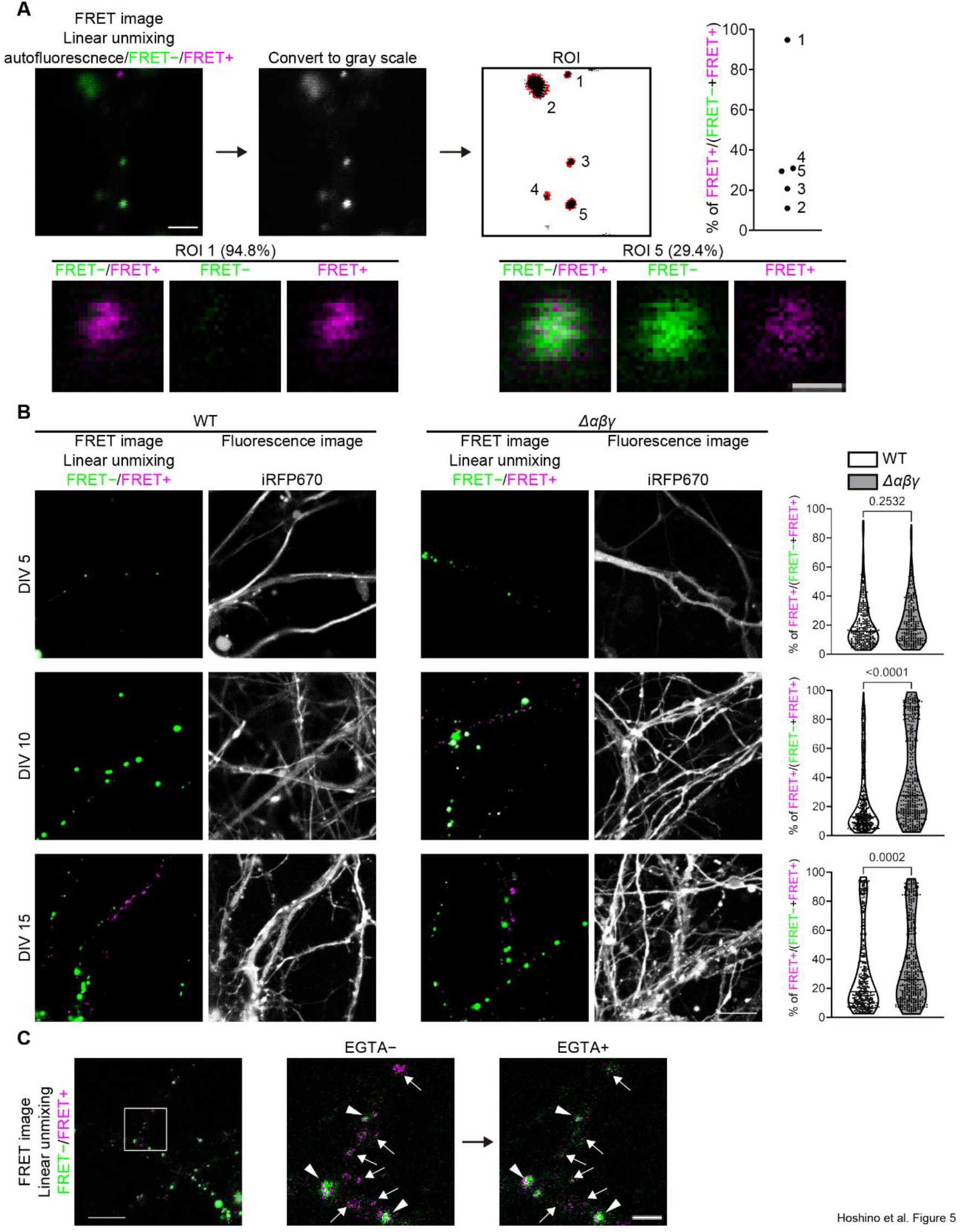
Endogenous Pcdhs inhibit the exogenous Pcdh homophilic interaction. (A) Process for analyzing % of FRET+ ratio in neurons. The acquired spectrum image is linear unmixed for autofluorescence, FRET− and FRET+. The image is converted to grayscale. A binary image was created, and ROIs were defined using the Fiji. % of fluorescent intensity for FRET+ relative to the sum fluorescent intensities of FRET− and FRET+ are plotted. ROI 1 and ROI 5 in the top panels were magnified in the bottom panels. Scale bar, 2 µm for the top and 0.5 µm for the bottom. (B) Cell morphologies and FRET observations in WT and Δαβγ neurons at DIV 5, 10, and 15 (left). Quantification showing the % of FRET+ intensity of each dot (n = 12 for each genotype and each DIV; line: median, dotted line: 25th/75th percentiles; Mann Whitney test) (right). Scale bar, 10 µm. (C) FRET+ dots disappeared upon EGTA addition. Arrows indicate FRET probes showing FRET+ signals and they disappear after EGTA addition. Arrowheads indicate FRET probes showing FRET− signals and they do not disappear after EGTA addition. Scale bar, 10 µm for the left panel, and 2 µm for the right panel. See also Figure S5.

Neurons were transfected with plasmids encoding the combined γB2FL-FRET probe and iRFP670 to visualize the cell morphology (Figures 5B and S5A). In DIV 5 cultures, almost no FRET signals in either WT neurons or Δαβγ were observed, resulting in a unimodal FRET+ ratio. In the DIV 15 cultures, some FRET probes showed FRET signals, resulting in a bimodal FRET+ ratio distribution. These results suggested that the homophilic interaction of Pcdh increases during development. Interestingly, in DIV 10 cultures, WT neurons showed a unimodal FRET+ ratio, as observed in DIV5 but Δαβγ neurons showed a bimodal FRET+ ratio, as observed in DIV 15. At each time point, γB2-FRET signals were observed more frequently in Δαβγ neurons than in WT neurons (Figure 5B). This indicated that endogenous Pcdh isoforms inhibited the homophilic interaction of the combined γB2-FRET probes.

To determine whether the FRET+ signals found in the neuron culture were calcium-dependent, we added EGTA to the medium. The γB2 protein showing FRET+ signals disappeared rather than showing FRET− signals, while those showing FRET− signals were still localized after EGTA addition (Figure 5C). The addition of EGTA to K562 cells expressing the combined γB2ΔICD-FRET probe also showed a disappearing reaction when the γB2ΔICD proteins did not accumulate strongly at the contact sites (Figures S5B and S5C). When the combined γB2ΔICD-FRET probe was transfected into neuron cultures, most of the γB2ΔICD proteins were localized as dots; however, some were localized as lines, likely observed at cell contact sites using K562 cells. The dotted γB2ΔICD proteins in neurons showed FRET− and FRET+ signals, but all the lined ones showed FRET+ signals (Figure S5C). In the lined localization, the combined γB2ΔICD-FRET probe localization was maintained after EGTA, and the FRET+ signals turned into FRET− signals (Figure S5C). However, the FRET+ signals of the dotted localization disappeared immediately after the addition of EGTA. This disappearance was also observed at the weak cell contact sites of K562 cells (Figure S5B). These results suggest that some γB2ΔICD proteins at the plasma membranes move from the cell contact sites; however, accumulated proteins that strongly interact at cell contact sites remain at the site after EGTA addition. Both the combined γB2FL-FRET probe and combined γB2ΔICD-FRET probe that show FRET− signals do not disappear and retain FRET− signals after EGTA addition. These results suggest that the γB2 proteins showing FRET− signals are protected by diffusion, although the γB2 proteins showing FRET+ signals are fragile after the homophilic interaction is interrupted by EGTA.

### Homophilically interacting γB2 proteins are rare at inhibitory synapses or excitatory synapses

Previous studies have shown that Pcdhγ localizes to subsets of synapses in cultured hippocampal neurons (24, 25). However, it remains unclear whether the synaptic Pcdh proteins interact homophilically. To investigate the relationships between synaptic localization and the state of the Pcdh homophilic interaction, we transfected plasmids to visualize synapses together with plasmids containing the combined γB2FL-FRET probe and iRFP670 into cultured hippocampal neurons. The co-transfection strategy for visualizing synapses is effective since the fluorescently labeled synaptic marker is of a neuron that shows the state of Pcdh homophilic interaction rather than the synaptic marker of the surrounding non-transfected neurons. FingR (GPHN)-Cherry was used to visualize gephyrin, an inhibitory synaptic marker (29, 30), while PSD95-TagRFP was used to visualize PSD95, an excitatory synaptic marker.

We prepared cultured hippocampal neurons from WT mice and observed FRET and synaptic markers on DIV 13-17 (Figures 6A, 6C, S6A, and S6B). We classified each γB2 dot based on whether they did or did not colocalize with synaptic markers. Most of the γB2 dots did not colocalize with synaptic markers, and showed both a low % of FRET+ and high % of FRET+ signals, likely in the observation without transfection of synaptic markers at DIV 15 (Figures 5B, 6B, and 6D). However, γB2 dots colocalizing synaptic markers showed a significantly low % of FRET+ signals for gephyrin and PSD95. These results suggest that Pcdhγ proteins at the inhibitory and excitatory synapses rarely interact homophilically.

**Figure 6.**
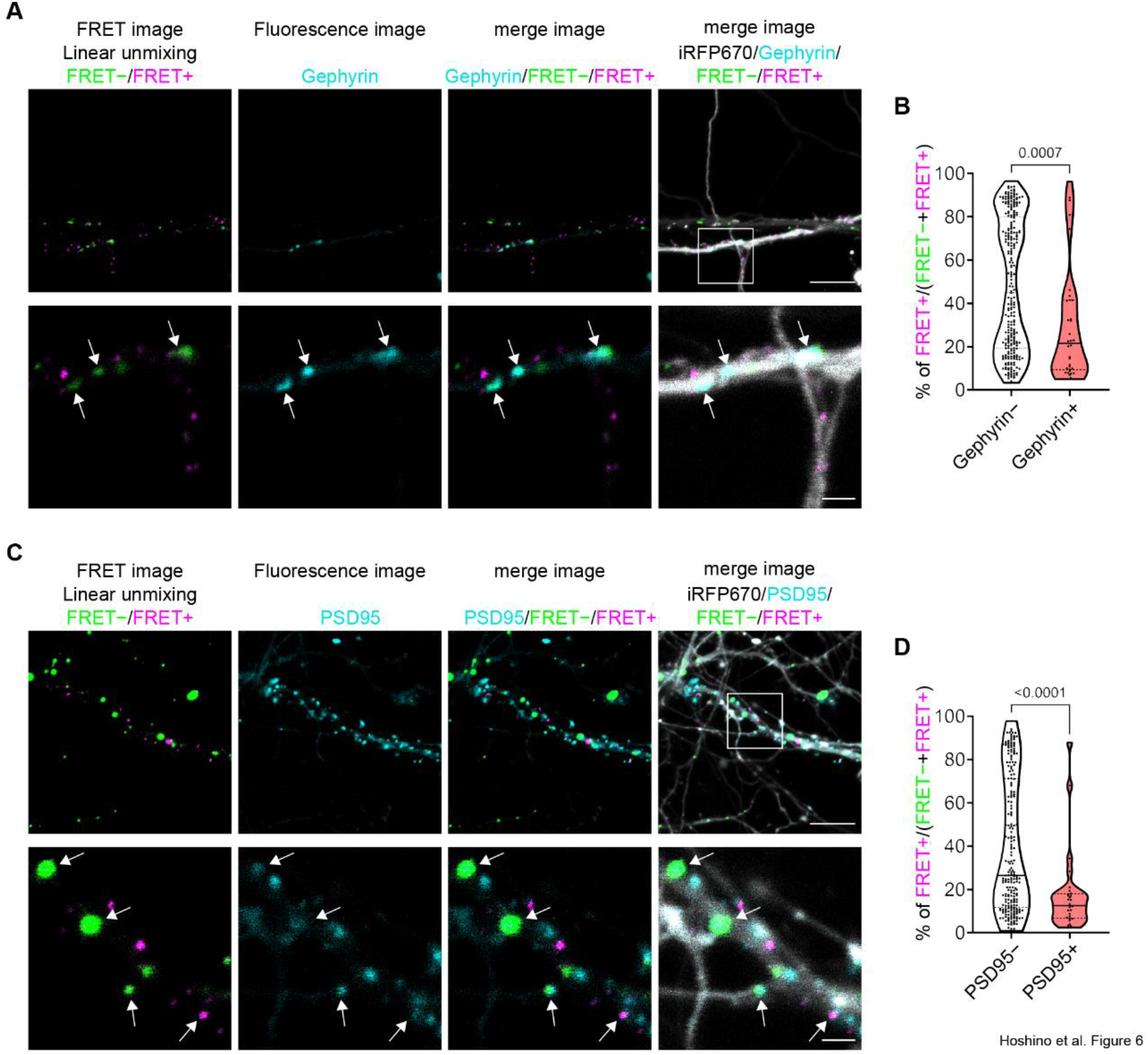
Pcdh at synapses are not homophilically interacting. (A) Linear unmixed images, FingR (GPHN)-Cherry, and iRFP670 are shown. The bottom panels are magnified images for the white square of the top panels. Arrows indicate the dots of the combined γB2FL-FRET probe colocalizing with gephyrin. Scale bar, 10 µm for the top and 2 µm for the bottom. (B) Quantification showing the % of FRET+ intensity of each dot (n = 11; line: median, dotted line: 25th/75th percentiles; Mann Whitney test) (C) Linear unmixed images, PSD95-TagRFP and iRFP670 are shown. The bottom panels are magnified images for the white square of the top panels. Arrows indicate the dots of the combined γB2FL-FRET probe colocalizing with PSD95. Scale bar, 10 µm for the top and 2 µm for the bottom. (D) Quantification showing the % of FRET+ intensity of each dot (n = 9; line: median, dotted line: 25th/75th percentiles; Mann Whitney test)

## Discussion

We visualized the homophilic Pcdh interaction in neurons. Previously, the Pcdh homophilic interaction was analyzed using a cell aggregation assay with K562 cells lacking endogenous cell adhesion molecules. However, observing where and when the interaction occurred in neurons was impossible.

The separated γB2-FRET probe could not be applied to visualize the Pcdh homophilic interactions in self-neurites (22) (Figures 1A, 1C, and 1E). The combined γB2FL-FRET probe solved the problem (Figures 1D, 1E, and 3A).

Acceptor photobleaching showed that the separated γB2ΔICD-FRET probe had a higher FRET efficiency than the combined γB2ΔICD-FRET probe (22) (Figure 1I). This may be due to a slight conformational change in the extracellular domains of γB2 or steric hindrance of CFP and YFP. However, the K562 cell aggregation assay showed that this influence did not interfere with the homophilic interaction between γB2 (Figure 1J). Additionally, similar to the native γB2 protein, the γB2FL-FRET probe showed *cis* interaction with the α4 isoform (Figure S1E) and could translocate it to the plasma membrane (Figure S1D).

One of the difficulties with ratio imaging using intermolecular FRET probes is the unequal distribution of CFP- and YFP-fused proteins because YFP can be directly excited by a laser used for CFP excitation. In the YFP-rich region, non-negligible fluorescence from YFP directly excited by a CFP-exciting laser was detected in the FRET channel (YFP emission with CFP excitation), which can cause a pseudo-positive FRET signal. While the separated γB2-FRET probes can be analyzed using ratio imaging with a 405 nm laser (Figure S2), which directly excites only 0.3% of YFP when compared to that at its optimal excitation wavelength (514 nm) (31), such a short-wavelength laser is phototoxic to neurons. To avoid phototoxicity, we used a 458 nm laser for CFP excitation. In this case, 10% of the YFP, compared to that at its optimal excitation wavelength, is directly excited, regardless of FRET. Therefore, ratio imaging of general intramolecular FRET probes and the separated γB2 FRET probe must be corrected with the expression levels of CFP and YFP (31, 32), or the FRET signals should be observed using fluorescence lifetime imaging microscopy (FLIM) or acceptor photobleaching. However, these methods are technically difficult, expensive, or irreversible to perform. Using the combined γB2-FRET probe, CFP and YFP were equally localized anywhere, because the CFP and YFP were fused to a single Pcdh protein, which we term as the direct excitation auto-normalizing intermolecular (DENIM) FRET probe.

To generate a DENIM FRET probe for visualizing homophilic protein-protein interactions, donor and acceptor fluorescent proteins must be fused at different places on amino acid sequences. In the γB2-FRET probe, CFP was fused to extracellular domain 1, and YFP was fused to extracellular domain 5. The FRET probe development can be started from the separated probe, and the donor and acceptor fluorescent proteins can be combined into a single protein molecule. DENIM FRET probes can be generated to visualize heterophilic interactions by fusing additional fluorescent proteins. When protein A is fused with CFP and protein B is fused with YFP to produce FRET signals, YFP can be fused with protein A and CFP can be fused with protein B to normalize the direct excitation of YFP. It should be confirmed that these additional fluorescent proteins do not show unintentional FRET signals and that they do not interfere with the interaction between proteins A and B.

To date, Pcdh homophilic interactions have been studied using a biochemical approach based on crystal structure, analytical ultracentrifugation, and surface plasmon resonance (3, 13–16, 33).

From the cell biology approach, cell aggregation assay using K562 cells has clearly shown their strict homophilic interaction (3, 4, 12). However, these methods do not provide detailed spatiotemporal information about neurons.

Pcdh homophilic interactions in neurons have been studied by transfecting matched or mismatched Pcdh isoforms (20, 21). They showed that Pcdh proteins from two different neurons colocalized more often in the matched isoform and that dendrite complexity in the cortex is regulated non-cell autonomously, which strongly suggests that Pcdh homophilic interaction occurs in neurons. Here, we first observed the Pcdh homophilic interaction in neurons using time-lapse imaging and succeeded in studying when and where the Pcdh interaction occurs in neurons.

The combined γB2FL-FRET probe was localized to the soma and neurites. As reported in a previous study (23), the combined γB2FL-FRET probe accumulated at neurite contacts, and FRET signals were observed at these sites. The γB2 proteins in the soma did not show FRET signals, but FRET signals were observed in the roots of neurites from the soma (Figure 3A). This suggests that the Pcdh homophilic interaction is involved in neurite growth or proper development of neurites.

Although Pcdhγ is highly expressed in the hippocampus and localizes at synapses, previous studies using cultured hippocampal neurons showed that Pcdhαβγ or Pcdhγ deletion did not affect the number of PSD95 and Gephyrin (20, 26). Our results show that γB2 proteins at synapses rarely interact homophilically, indicating that Pcdhγ regulates synaptic specificity rather than the total synaptic number. In contrast, in cortical neurons, Pcdhγ deletion increased the number of synapses, and the interaction of Pcdhγ with Neuroligin-1 and Neuroligin-2 is involved in this process (34, 35). Such brain region-or cell type-specific functions and molecular mechanisms remain unknown.

In this study, we first demonstrated the relationship between Pcdh homophilic interactions and synaptic regulation in cultured hippocampal neurons.

There are still many questions regarding the homophilic interaction of Pcdh, which is unique to the cell surface recognition molecule utilizing stochastic combinatorial expression of over 50 isoforms. In this study, we demonstrated colocalization analysis using the γB2-FRET probe and synaptic marker. To understand molecular mechanism of the Pcdh homophilic interaction for synaptic regulation, self-avoidance, or any other functions, we can analyze the spatiotemporal information of Pcdh homophilic interaction using the γB2-FRET probe along with localization of other molecules or the information during biological events. The γB2-FRET probe opens the door to investigating the role of Pcdh homophilic interactions in the CNS.

## Materials and Methods

Detailed information on materials and methods used in this study is found in the SI Appendix. Further information and requests for resources and reagents should be directed to and will be fulfilled by Takeshi Yagi.

### FRET observation with ratio imaging

CFP was excited with a 405 nm or 458 nm laser using LSM780 (Zeiss) with a Plan-Apochromat ×63, 1.40 NA oil immersion objective lens. Using either laser, the donor- and acceptor-emitted light was collected at 463–500 nm and 520–620 nm, respectively. The ratio analysis was performed using cellSens (Olympus).

### FRET observation with spectrum imaging followed by linear unmixing

CFP was excited with a 458 nm laser using LSM780 (Zeiss) with a Plan-Apochromat ×63, 1.40 NA oil immersion objective lens, and the emitted light was collected over a spectrum with a width of 8.9 nm. Spectrum imaging was performed to collect the reference spectrum, cultured hippocampal neurons for autofluorescence, intracellular space of K562 cells expressing the combined γB2ΔICD-FRET probe for FRET−, and cell contact site of K562 cells expressing the combined γB2ΔICD-FRET probe for FRET+. The fluorescence of the sample was measured using spectral imaging, and linear unmixing was performed using Zen (Zeiss). The unmixed image was converted to binary and we obtained ROIs for each dot of the combined γB2FL-FRET probes using Fiji. The autofluorescence, FRET−, and FRET+ intensities were measured for each ROI using the Fiji software. The ratios of the three components were calculated using Excel software. ROIs that contained more than 10% autofluorescence were excluded from FRET analysis. Nine out of the 1856 ROIs were excluded. Colocalization analysis using synaptic markers was manually performed. The ROI for each dot of the combined γB2FL-FRET probes was classified as non-colocalized or colocalized with synaptic markers.

## Acknowledgments

We would like to thank Dr. Masayoshi Mishina for providing the EF1a Flp mice. This work was supported by JST PRESTO Program (No. JPMJPR2045) to T.K., JSPS KAKENHI Grant Number JP16H06280, Grant-in-Aid for Scientific Research on Innovative Areas ― Platforms for Advanced Technologies and Research Resources “Advanced Bioimaging Support” to E.T., Grants-in-Aid for Transformative Research Areas (A) (20H05899) to E.T., the MEXT Grant-in-Aid for Scientific Research (A) from JSPS (No. 18H04016) to T.Y., Grant-in-Aid for Scientific Research on Innovative Areas “Integrated analysis and regulation of cellular diversity” (No. 20H05035) to T.Y., for Scientific Research on Transformative Research Areas (A) Adaptive Circuit Census (No. 22H05498) to T.Y., National Institutes of Health Grants R01 MH117790 to T.Y., and in part by the Planned Collaborative Project and the Cooperative Study Program of the National Institute for Physiological Sciences, Japan to T.Y.

