## Supplemental Information for "Visualization of *trans* homophilic interaction of clustered protocadherin in neurons"

[a]KOKORO-Biology Group, Graduate School of Frontier Biosciences, Osaka University, Suita, Osaka 565-0871, Japan. [b]Department of Biomolecular Science and Engineering, SANKEN (The Institute of Scientific and Industrial Research), Osaka University, Ibaraki, Osaka 567-0047, Japan. [c]Japan Science and Technology Agency (JST), Precursory Research for Embryonic Science and Technology (PRESTO), Kawaguchi, Saitama 332-0012, Japan. [d]Department of Biochemistry and Cellular Biology, National Institute of Neuroscience, National Center of Neurology and Psychiatry, Tokyo 142-8501, Japan. [e]Clinical Medicine Research Laboratory, Shonan University of Medical Sciences, Yokohama 244-0806, Japan. [f]Department of Anatomy, Faculty of Medicine, Hokkaido University, Sapporo, Hokkaido 060-8638, Japan.

*Corresponding author: Takeshi Yagi

**This file includes:**

- Supporting text
- Figures S1 to S6
- Tables S1
- SI References

### SI Appendix – Material

#### Material availability

The Pcdhγ floxed mice generated in this study were deposited in RIKEN BRC (https://web.brc.riken.jp/en/) (RBRC02800). The PcdhγB2-FRET mice generated in this study were deposited in RIKEN BRC (https://web.brc.riken.jp/en/) (RBRC11902).

#### Mice studies

Pcdhγ-floxed mice and γB2-FRET mice were generated as described in the Methods Details section. Embryonic male and female mice were used for cultured hippocampal neurons, and postnatal male and female mice were used for the evaluation of self-crossings and qPCR and western blotting.
All animal care protocols and experiments were performed according to guidelines approved by the Animal Experiment Committee of the Graduate School of Frontier Biosciences at Osaka University (approval number: FBS-22-008) and the Animal Care and Use Committee of the National Institute of Neuroscience, National Center of Neurology and Psychiatry (approval number: 2020007). Mice were housed in groups under a 12 h light/12 h dark cycle.

#### Primary hippocampus culture

The media used in this study were MEM (Gibco) or FluoroBrite DMEM (Gibco) supplemented with 5.5% FBS (Invitrogen), 2% B27 supplement (Gibco), 1 mM GlutaMAX (Gibco), 100 units/mL Penicillin-100 µg/mL streptomycin (Gibco), and 10 mM HEPES (1 M) (Gibco). The hippocampi from E16–E18 mice were dissected in HBSS (Gibco) with 10 mM HEPES (1 M) (Gibco). The hippocampi were treated with Neuron Dissociation Solutions (Wako) according to the manufacturer's protocol. The dissociated cells were washed in a MEM-based medium. The cells were then electroporated using an Amaxa 4D-Nucleofector (Lonza). The transfected cells were suspended in a MEM-based medium and plated on a poly L-lysine-treated CELLview glass bottom dish (Advanced TC, 4-compartments) (Greiner). After 2 h, the medium was replaced with the FluoroBrite DMEM-based medium. The following day, 5 mM cytosine β-D-arabinofuranoside hydrochloride (Sigma-Aldrich) was added to a final concentration of 5 nM. The cells were incubated at 37°C in humidified air containing 5% $CO_2$, without any medium changes. WT mice used in cultured hippocampal neurons are littermates of Δαβγ with Taf7 transgene mice and the WT mice have Taf7 transgene.

#### K562 culture

K562 cells (RIKEN BRC) from female chronic myelogenous leukemia patients in blast crisis were maintained in Iscove's modified Dulbecco's medium (IMDM, Wako Pure Chemical Industries) supplemented with 9% (v/v) FBS (Gibco) and 45 µg/mL kanamycin at 37°C in humidified air containing 5% $CO_2$. K562 cells were electroporated using an Amaxa 4D-Nucleofector (Lonza).

#### HEK293T culture

HEK293T cells (RIKEN BRC) from female human embryonic kidneys were maintained in Dulbecco's modified Eagle's medium (DMEM, Sigma-Aldrich) supplemented with 10% (v/v) fetal bovine serum (FBS, Biowest) at 37°C in humidified air containing 5% $CO_2$. HEK 293T cells were transfected with polyethyleneimine "MAX" (Cosmo Bio).

#### Plasmid construction

All expression vectors for Pcdh were subcloned into pCX vectors. The combined γB2FL-FRET probe and the combined γB2ΔICD-FRET probe were generated using the separated γB2FL-FRET probes and the separated γB2ΔICD-FRET probe by enzyme digestion and ligation. α4ΔICD-mCherry was generated by the deletion of ICD (V745–Q947) of α4, followed by NheI site and mCherry. The expression vector for mCherry was a kind gift from Hiroaki Kobayashi, Osaka University used in Yanagida et al(1). The expression vector for iRFP670 was subcloned from piRFP670-N1 (Addgene #45457) into the pCX vector. FingR (GPHN)-Cherry(2) was modified from pCAG_GPHN.FingR-eGFP-CCR5TC (Addgene #46296).

**Antibodies**
A polyclonal antibody against the Pcdhγ constant region (PcdhγCR) was raised in guinea pigs and rabbits by using Pcdhγ-A12 (NM_033595.4, aa 809–932). To express the glutathione S-transferase fusion proteins, we subcloned the corresponding cDNA fragments into the pGEX4T-2 plasmid (GE Healthcare, Buckinghamshire, UK). Immunization and affinity purification were performed as previously reported (3, 4).

### SI Appendix – Methods

#### Generation of Pcdhγ floxed mice

To introduce loxP sequences both upstream and downstream of the Pcdhγ CR1 exon (γCR1), a targeting vector was constructed using bacterial artificial chromosome (BAC) modification methods and the Red recombination system (5). The modifications were performed by transferring purified mouse BAC RP23-332K20 into the E. coli strain EL350 via electroporation. Three genomic DNA fragments, a fragment containing a loxP sequence and γCR1, and two homology fragments for BAC modification, were PCR-amplified using mouse BAC as the template, and the gCR1A-F and gCR1A-R, gCR1B-F, and gCR1B-R, and gCR1C-F and gCR1C-R primer pairs, respectively. These DNA fragments were subcloned into the *pBTloxP2* plasmid, which contained a *loxP* and *frt*-flanked neomycin resistance (*neo*$^r$) cassette. The *neo*$^r$ gene with *floxed-γCR1* and homology arms were excised by *Kpn*I/*Sac*I digestion and gel-purified. The purified DNA fragment was electroporated into EL350 cells containing BAC RP23-332K20, which had been induced for Red recombination by prior growth at 42°C for 15 min. The transformants were selected on kanamycin-containing plates. Homology fragments for retrieval were amplified using the gCR1D-F, gCR1D-R, gCR1E-F, and gCR1D-R primer pairs and subcloned into *pBRSDT*, which contains the gene for diphtheria toxin A, between the *Hind*III and *Nhe*I restriction sites. We then retrieved the BAC DNA fragments that contained the *floxed-γCR1* and *frt*-flanked *neo*$^r$ cassette and inserted them into *pBRSDT* using Red recombination. The retrieved plasmid was used as a target vector.
The linearized targeting vector was electroporated into TT2 ES cells, and homologous recombinant ES clones were screened by Southern hybridization with a probe isolated by PCR using mouse BAC as a template and ProbeA-F and ProbeA-R, and ProbeB-F and ProbeB-R primer pairs. Recombinant ES cell clones were injected into ICR mouse blastocysts and male chimeras were bred with C57BL/6J mice. Finally, the *neo*$^r$ cassette was removed from *γCR1-floxed neo* mice by crossing them with EF1a Flp mice (6).

#### Generation of γB2-FRET mice

To induce conditional overexpression of the combined γB2FL-FRET probe, its coding sequences were cloned into the Cre-dependent expression plasmid vector, which also harbored appropriate homology arms for recombineering to the mouse ROSA26 locus containing BAC (7), via *Pac*I and *Pme*I digestion and T4 ligase-mediated ligation. The resulting plasmid was digested with *Sal*I and *Xho*I and then electroporated into *E. coli* EL350 containing BAC (8). The appropriately recombined BAC clone was selected and transferred into ElectroMAX DH10B and then purified with NucleoBond BAC100 (Macherey-Nagel, 740579). BAC DNA was linearized at a *Fse*I site on the backbone vector and microinjected into fertilized eggs, collected from super-ovulated B6C3F1 (SLC, Japan) females, at a concentration of 0.5 ng/μL. The eggs were transferred into the oviducts of surrogate mothers to raise pups. Genomic DNAs were prepared from the tail of the newborn, and transgenic mice were screened by PCR using the pCX1624-F1 and FRET-R primer pairs to confirm intact insertion of the combined γB2FL-FRET probe. To further ensure full-length integration of the ROSA locus containing BAC in transgenic founders, the primer set (F/pBAC-4 and R/pBAC-3, RP23-244D9 end F1, and pBAC-R2) was used to amplify the flanking region of *Fse*I site on the backbone vector. Three transgenic founders were obtained, and the combined γB2FL-FRET probe expression levels were quantified by qRT-PCR using the primer set (B2m-real-F, B2m-real-R, Venus-gB2-F, and FRET-EC5-qPCR-R). The mouse line with the highest expression level was selected for use in this study.

**Cell aggregation assay**
The transfected K562 cells were rotated at 30 rpm overnight at 37°C in a humidified atmosphere containing 5% $CO_2$. Before imaging, the cells were decanted into glass-bottomed dishes coated with 0.1% polyethyleneimine. The images were obtained using an LSM780 (Zeiss) confocal microscope equipped with a Plan-Apochromat ×10, 0.45 NA objective lens and a Plan-Apochromat ×63, 1.40 NA oil immersion objective lens.

**Acceptor photobleaching**
CFP and YFP were excited using 405 nm and 515 nm lasers, and their emissions were collected using bandpass filters set at 460–500 nm and 530–630 nm, respectively, by FV-1000 confocal microscopy using an IX81 microscope equipped with a ×60, 1.35 NA oil-immersion objective lens (UPLSAPO60XO) (Olympus). Fluorescence intensities were measured using the FV10-ASW software (Olympus). FRET efficiency was calculated using the following formula: FRET efficiency = $1- (F_{BB}/F_{AB})$, where $F_{BB}$ and $F_{AB}$ are the fluorescence intensities of CFP before and after photobleaching, respectively.

**EGTA addition in K562 cells and cultured hippocampal neurons**
To adjust the osmotic pressure, 500 mM EGTA dissolved in water, water, and the medium was mixed at a ratio of 1:5:4 in volume for the K562 cell culture and 1:0:9 for the neuron culture. The resultant solution was added manually to a glass-bottom dish at a volume of 10% of the medium to reach a final concentration of 5 mM EGTA during time-lapse imaging of 1 s/frame.

**qRT-PCR**
Genotypes with $PV^{PV\text{-}Cre/WT}$; $Pcdh\gamma^{flox/flox}$; $\gamma B2\text{-}FRET^{Tg/WT}$; $Ai14^{KI/WT}$ were used. P24 mice were anesthetized and the cerebellums were dissected in ice-cold HBSS. The cerebellum was immediately frozen at −80°C. mRNA was acquired using RNeasy (Qiagen), according to the manufacturer’s protocol. Briefly, the cerebellum was submerged in ice-cold RLT Buffer and homogenized with ShakeMan2 (BMS) at room temperature. The extract was treated with DNase, and total mRNA was reverse-transcribed using a PrimeScript RT reagent Kit (Perfect Real Time) (Takara). The amount of cDNA was quantified using TB Green Premix Ex Taq™ II (Tli RNaseH Plus) (Takara), and qPCR was performed using the StepOnePlus Real-Time PCR System (Thermo Fisher).

**Immunoprecipitation**
K562 cells were electroporated using an Amaxa 4D-Nucleofector (Lonza). After 18 h, cells were collected and washed with ice-cold PBS. The cells were treated with lysis buffer (25 mM Tris-HCl (pH 7.4), 50 mM NaCl, 25 mM NaF, 1 mM EDTA, 1% Nonidet-P40, 5% glycerol) supplemented with a cOmplete Protease Inhibitor Cocktail (Roche). The amount of protein in the buffer was quantified by the BCA method (Thermo Fisher) using a microplate reader (Bio-Rad). Next, 500 µg of protein was diluted in 500 µL of lysis buffer. Samples (5 µL and 0.5 µL) were acquired from this diluted solution for the input sample. Protein G Sepharose 4 Fast Flow (GE Healthcare) was washed and resuspended in PBS. The sample (494.5 µL) was then treated with the beads to exclude the proteins that were non-specifically bound to the beads in a microtube rotator at 4°C for 1 h. The supernatant was then transferred into a new tube. The antibody (Rabbit polyclonal anti-Pcdhγ CR) was added to the newly washed beads and rotated at 4°C for 1 h, and the beads were washed with PBS. The beads with antibodies were then applied to the samples and rotated at 4°C for 2 h. The beads were then washed with lysis buffer and PBS. The IP sample and input were treated with the sample buffer (50 mM Tris-HCl, pH 7.5, 2% sodium dodecyl sulfate, 4% glycerol, 2% 2-Mercaptoethanol, and 0.02% bromophenol blue) at 95°C for 15 min.

**Western blot**
P25 mice were anesthetized and the cerebellums were dissected in ice-cold HBSS. The cerebellum was homogenized with ShakeMan2 (BMS) in homogenization buffer (50 mM Tris-HCl pH7.5, 150 mM NaCl, and 1 mM EDTA), and Triton X-100 (Nacalai Tesque) was added to a final concentration of 10%. The brain lysates or co-IP K562 proteins were electrophoresed with Bullet PAGE One Precast Gel (Nacalai Tesque) in an electrophoresis buffer (25 mM Tris, 192 mM

glycine, and 0.1% sodium dodecyl sulfate), and the proteins were transferred onto a nitrocellulose membrane (Bio-Rad) in transfer buffer (25 mM Tris, 192 mM glycine, and 10% methanol). The membrane was washed in TBS-T (200 mM NaCl, 20 mM Tris, 0.1% polyoxyethylene sorbitan monolaurate (Tween-20)) and blocked using a blocking buffer (4% skim milk in TBS-T). The primary antibody (Guinea pig polyclonal anti-Pcdhγ CR, Mouse monoclonal anti-Pcdhα CR (4F8) (9) was diluted in the blocking buffer and incubated overnight at 4°C. The next day, the membrane was washed with TBS-T, and the HRP-conjugated secondary antibody was diluted in the blocking buffer and incubated at room temperature for 2 h. The membrane was washed with TBS-T and treated with Chemi-Lumi One (Nacalai Tesque). Data were obtained using ChemiDoc MP (Bio-Rad).

**Quantification of self-crossings in Purkinje cells**

The mice were anesthetized with isoflurane and perfused with ice-cold ACSF (126 mM NaCl, 3 mM KCl, 1.3 mM $MgSO_4$, 2.4 mM $CaCl_2$, 1.2 mM $NaHPO_4$, 26 mM $NaHCO_3$, and 10 mM glucose). The brain was cut using a VT1200S vibrating blade microtome (Leica) and allowed to recover at 33°C for 1 h. Whole-cell voltage-clamp recordings were performed at room temperature. To load biocytin into the Purkinje cells, whole-cell recordings were maintained for 40 min at −70 mV with glass pipettes (5–7 MΩ) filled with a recording solution (130 mM K-gluconate, 8 mM KCl, 1 mM $MgCl_2$, 600 nm EGTA, 10 mM HEPES, 3 mM MgATP, 500 nm $Na_2GTP$, 10 mM Na-phosphocreatine, and 0.2% biocytin; pH adjusted to 7.3 using KOH). Slices were fixed with 4% PFA overnight at 4°C. The next day, the slices were washed with PBS and incubated with streptavidin and Alexa Fluor 488 conjugates for 3–6 days at room temperature. To avoid compressing the slices with a cover glass and slide glass, vinyl tape was placed on the glass to make room for the slices. The labeled Purkinje cells were imaged using an LSM780 (Zeiss) with a Plan-Apochromat ×40, 1.4 NA oil immersion objective lens at 0.5 µm intervals for the z-stack images. The self-crossings were counted in a 70.05 × 70.05 µm region of interest (ROI). The ROI was determined as follows. First, a marker was pointed at the center of the cell body. Second, we identified a marker at the end of the dendrites furthest from the cell body. Third, the centers of the two markers (cell body and dendrite) were calculated, and the center was also the ROI center. The self-crossing points were counted manually using the CellCounter plugin in Fiji software.

**Immunohistochemistry**

Three genotypes at P24-25 of either sex were used for immunostaining: control mice ($PV^{WT/WT}$; $Pcdh\gamma^{flox/flox}$; $\gamma B2\text{-}FRET^{WT/WT}$; $Ai14^{WT/WT}$), Pcdhγ-cKO ($PV^{PV\text{-}Cre/WT}$; $Pcdh\gamma^{flox/flox}$; $\gamma B2\text{-}FRET^{WT/WT}$; $Ai14^{KI/WT}$), and Pcdhγ cKO + γB2-FRET cOE mice ($PV^{PV\text{-}Cre/WT}$; $Pcdh\gamma^{flox/flox}$; $\gamma B2\text{-}FRET^{Tg/Tg}$; $Ai14^{WT/WT}$). Mice were anesthetized with isoflurane and perfused with ice-cold PBS. The brain was quickly removed and cryo-protected with compound, followed by indirect freezing with liquid nitrogen. Sagittal slices (10 µm thick) were cut using a cryostat (CM3050 S, Leica) and mounted on slide glass. Slices were fixed with 3% glyoxal solution (pH5.0) for 20 min at room temperature (RT). After washing with PBS thrice for 10 min, the slices were blocked using a blocking solution (2.5% normal goat serum, 0.1% Triton X-100 and 0.25% λ-carrageenan in PBS) for 1 h at RT. Then the slices were incubated with primary antibodies in a blocking solution overnight at RT. Anti-Pcdhγ CR (final concentration: 4.6 µg/mL, guinea pig) and anti-GFP (final concentration: 2.1 µg/mL, rat) were used. After washing in PBS thrice for 10 min, the slices were incubated with secondary antibodies in PBS containing 0.1 % Triton X-100 for 2 h at RT. See below for diluted secondary antibodies. Alexa Flour 647 (1:1000, guinea pig, Invitrogen), Alexa Flour 488 (1:1000, rat, plus, Invitrogen). The slices were imaged using an LSM780 (Zeiss) with a Plan-Apochromat ×40, 1.4 NA oil immersion objective lens.

**Quantification and statistical analysis**

The bar graphs display data as described in each figure legend. Statistical tests were performed using GraphPad Prism. $n \geq 2$ experiments were performed for each experiment that was statistically analyzed. Significant differences are indicated by number for each analysis in the figures. A p value $< 0.05$ was considered significant.

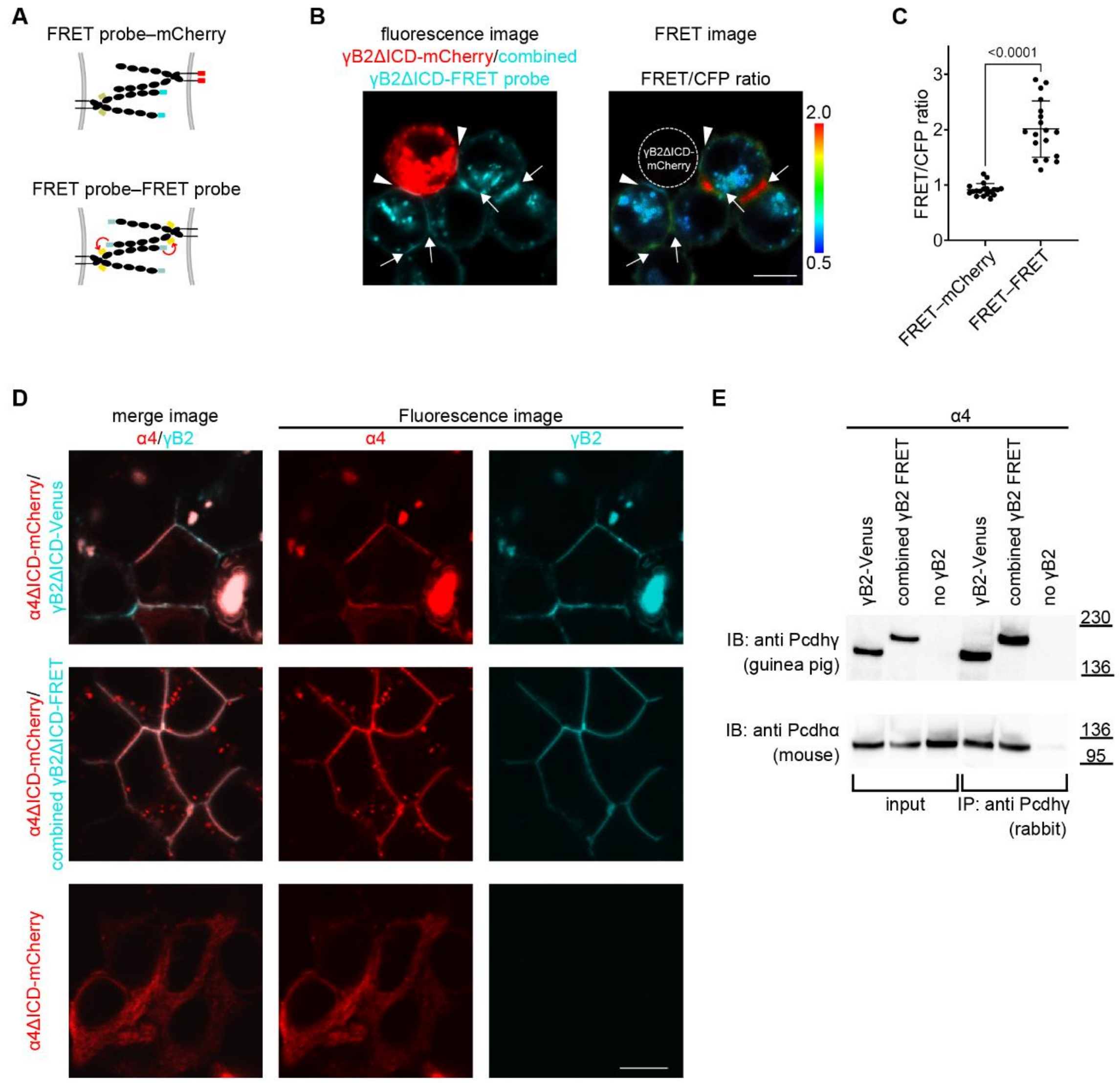


**Fig. S1. Combined γB2-FRET probe visualizes trans homophilic interaction and cis interaction with α4 is maintained, related to Figure 1.**

(A) Schematic of coculture of the combined γB2ΔICD-FRET probe-expressing cell and γB2ΔICD-mCherry-expressing cell. Red ovals represent mCherry. The cell contact sites between a combined γB2ΔICD-FRET probe-expressing cell and a γB2ΔICD-mCherry-expressing should not cause FRET if the FRET signal properly reflects the trans γB2 homophilic interaction.
(B) Ratio imaging of the coculture of the combined γB2ΔICD-FRET probe-expressing cells and γB2ΔICD-mCherry-expressing cells. Arrowheads indicate the cell contact sites between a combined γB2ΔICD-FRET probe-expressing cell and a γB2ΔICD-mCherry-expressing cell. Arrows indicate the cell contact sites between two combined γB2ΔICD-FRET probe-expressing cells. Scale bar, 10 µm.
(C) Quantification of FRET/CFP ratio of the cell contact sites of a combined γB2ΔICD-FRET probe-expressing cell and a γB2ΔICD-mCherry-expressing cell, and that of two combined γB2ΔICD-FRET probe-expressing cells (n = 9; mean ± SD; Welch’s t-test).
(D) Fluorescence image in HEK293T cells to observe the α4ΔICD-mCherry localization to plasma membrane delivered by γB2ΔICD-Venus or combined γB2ΔICD-FRET probe. Scale bar, 10 µm.
(E) Co-immunoprecipitation and Western blotting showing the cis interaction of γB2FL-Venus and the combined γB2FL-FRET probe with α4FL.

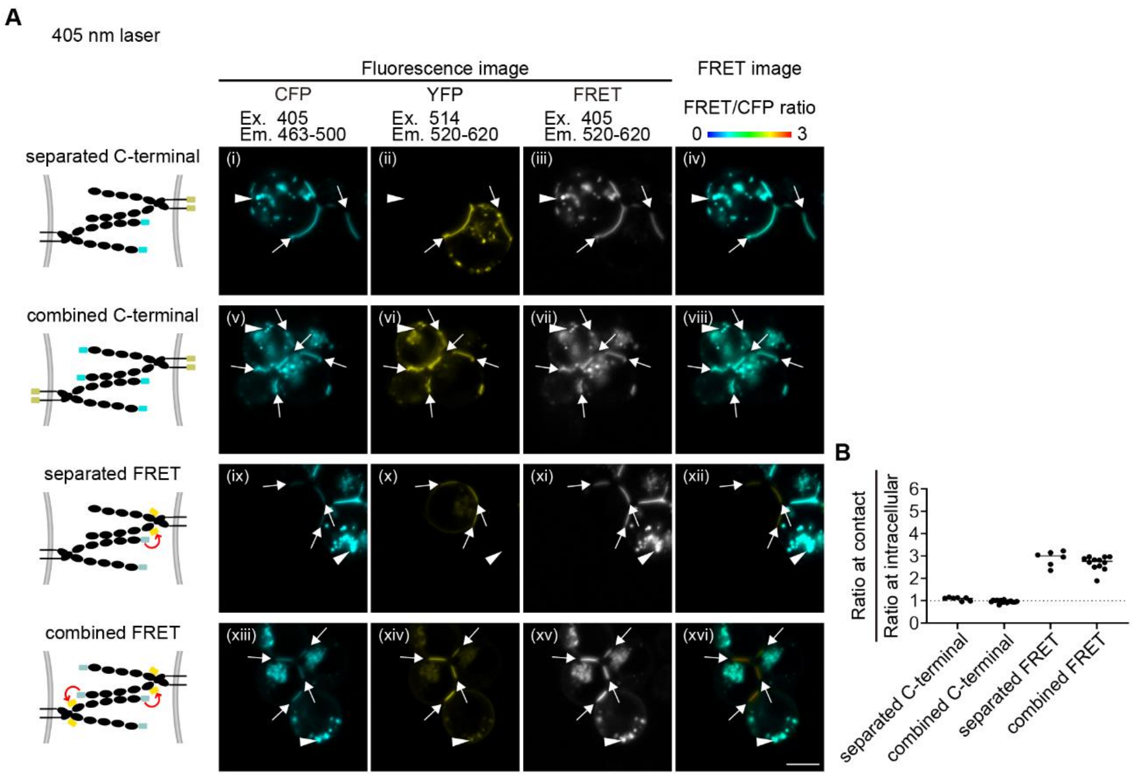


**Fig. S2. With a 405 nm laser, the separated and combined γB2-FRET probe can be used, related to Figure 2.**

(A) Ratio images of the separated and combined γB2ΔICD-C-terminal negative control and γB2ΔICD-FRET probes expressed in K562 cells with a 405 nm laser. Abbreviations: Em, emission; Ex, excitation. Arrows indicate the cell contact sites. Arrowheads indicate intracellular spaces. Scale bar, 10 µm.
(B) Quantification of the FRET/CFP ratio at cell contact sites normalized with the one at intracellular spaces (n = 3; mean).

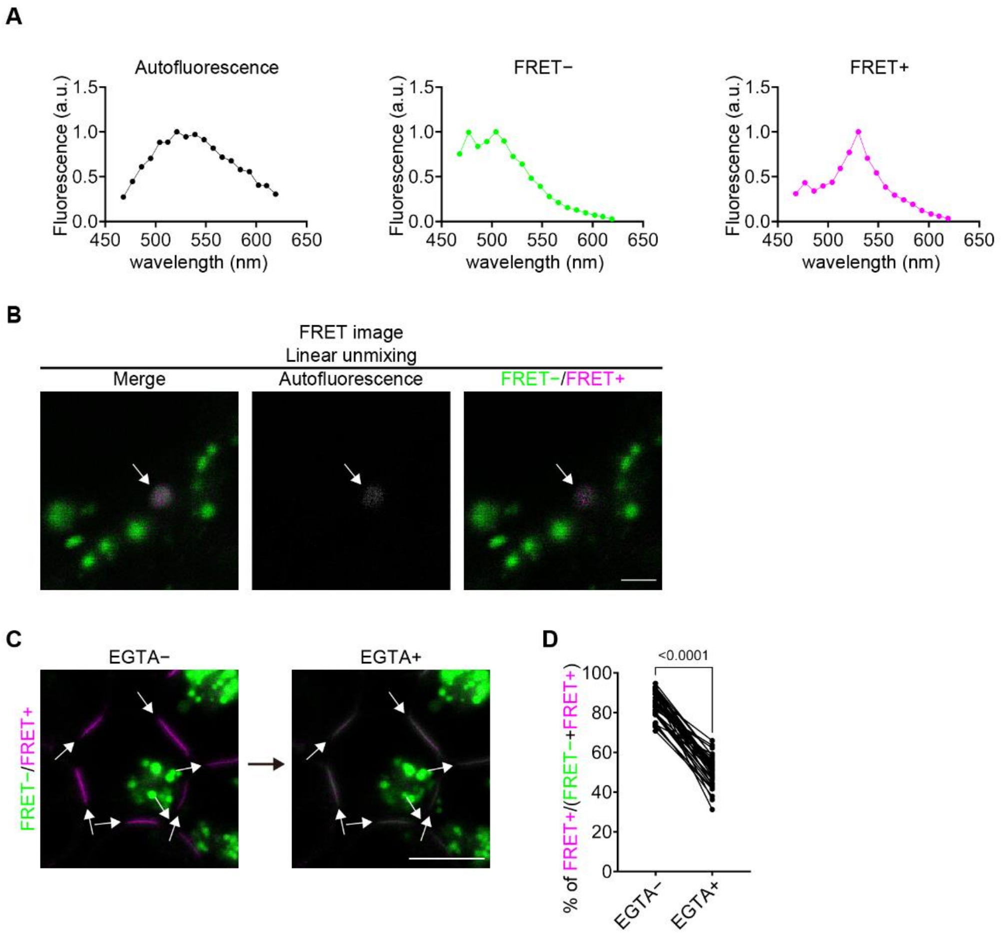


**Fig. S3. Spectrum imaging followed by linear unmixing is applicable, related to Figure 3.**

(A) Fluorescence spectra of autofluorescence, FRET−, and FRET+.
(B) Linear unmixed images of cultured neurons expressing the combined γB2-FRET probe. There were dots whose fluorescence signal was unmixed as autofluorescence (arrow). Such dots labeled as autofluorescence were 9 dots out of 1856 dots (See METHODS). Scale bar, 2 μm.
(C) Linear unmixed images of K562 cells expressing combined γB2ΔICD-FRET probe before and after EGTA addition. Arrows indicate cell contact sites. Scale bar, 10 μm.
(D) Quantification of the FRET signals before and after EGTA addition (n=7; mean ± SD; Wilcoxon matched-pairs rank test).

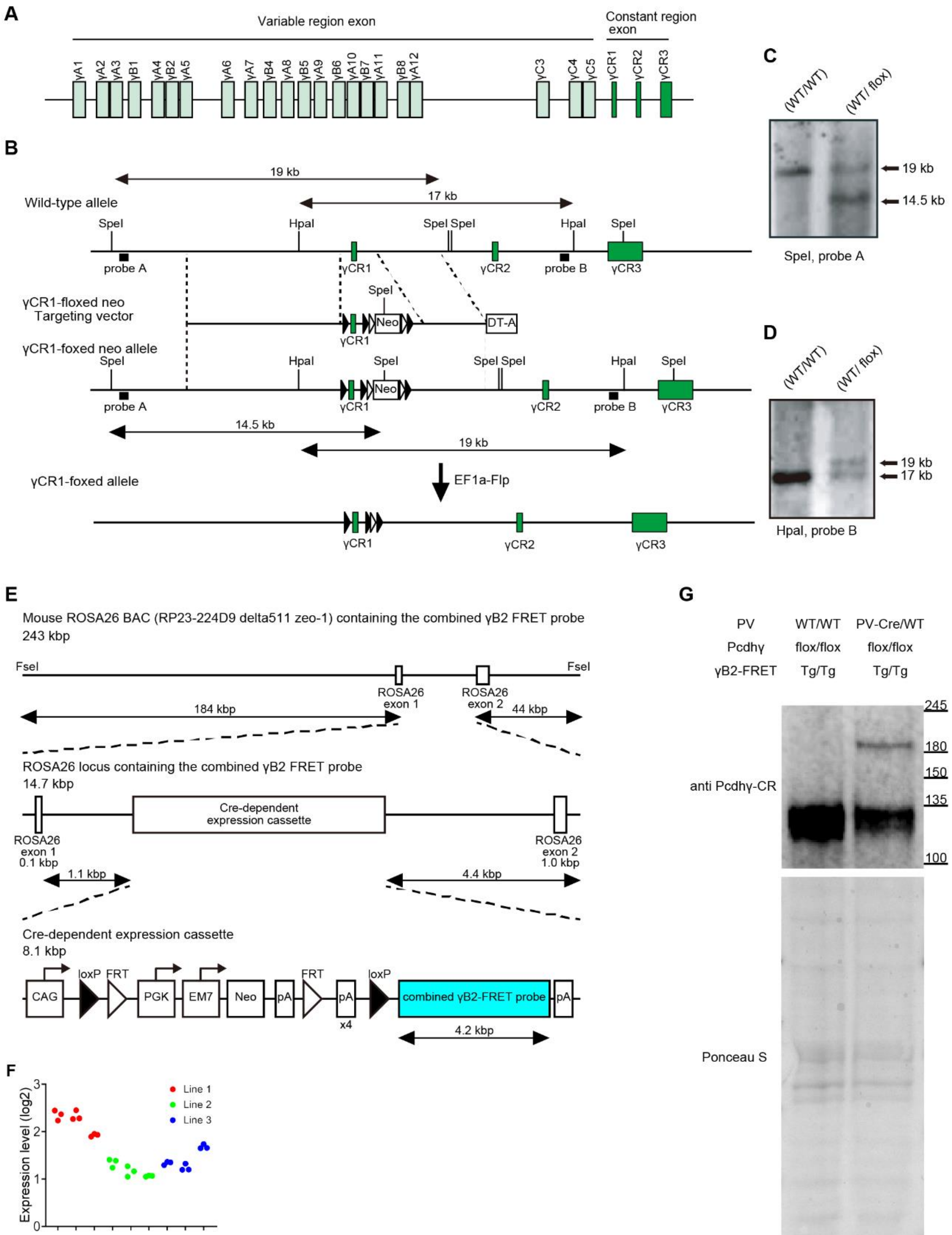


**Fig. S4. Generation of Pcdhγ-floxed mice and γB2-FRET mice, related to Figure 4.**

(A) Genomic structure of the *Pcdhg* genes.
(B) Schematic of the targeting constructs for the γCR-1floxed allele. The filled and open triangles represent *loxP* and *frt* sites, respectively.

(C) Southern blotting of genomic DNA isolated from homologous recombinant ES cells digested with *SpeI* using Probe A.
(D) Southern blotting of genomic DNA isolated from homologous recombinant ES cells digested with *SpeI* using Probe B.
(E) Schematic of the Cre-dependent combined γB2-FRET probe overexpression cassette recombined in a BAC.
(F) Expression levels quantified with qRT-PCR for three mouse lines generated in this study. The expression levels of the combined γB2-FRET probe were normalized with endogenous control, *β2-Microglobulin*. Red, green, and blue indicate each mouse line, and the analysis was performed three times for each mouse and three mice were analyzed for each mouse line.
(G) Western blotting showing Cre-dependent decreased expression levels of endogenous Pcdhγs (~ 120 kDa) and Cre-dependent overexpression of the combined γB2-FRET probe (~ 180 kDa) in Pcdhγ cKO + γB2-FRET cOE mice compared with control mice.

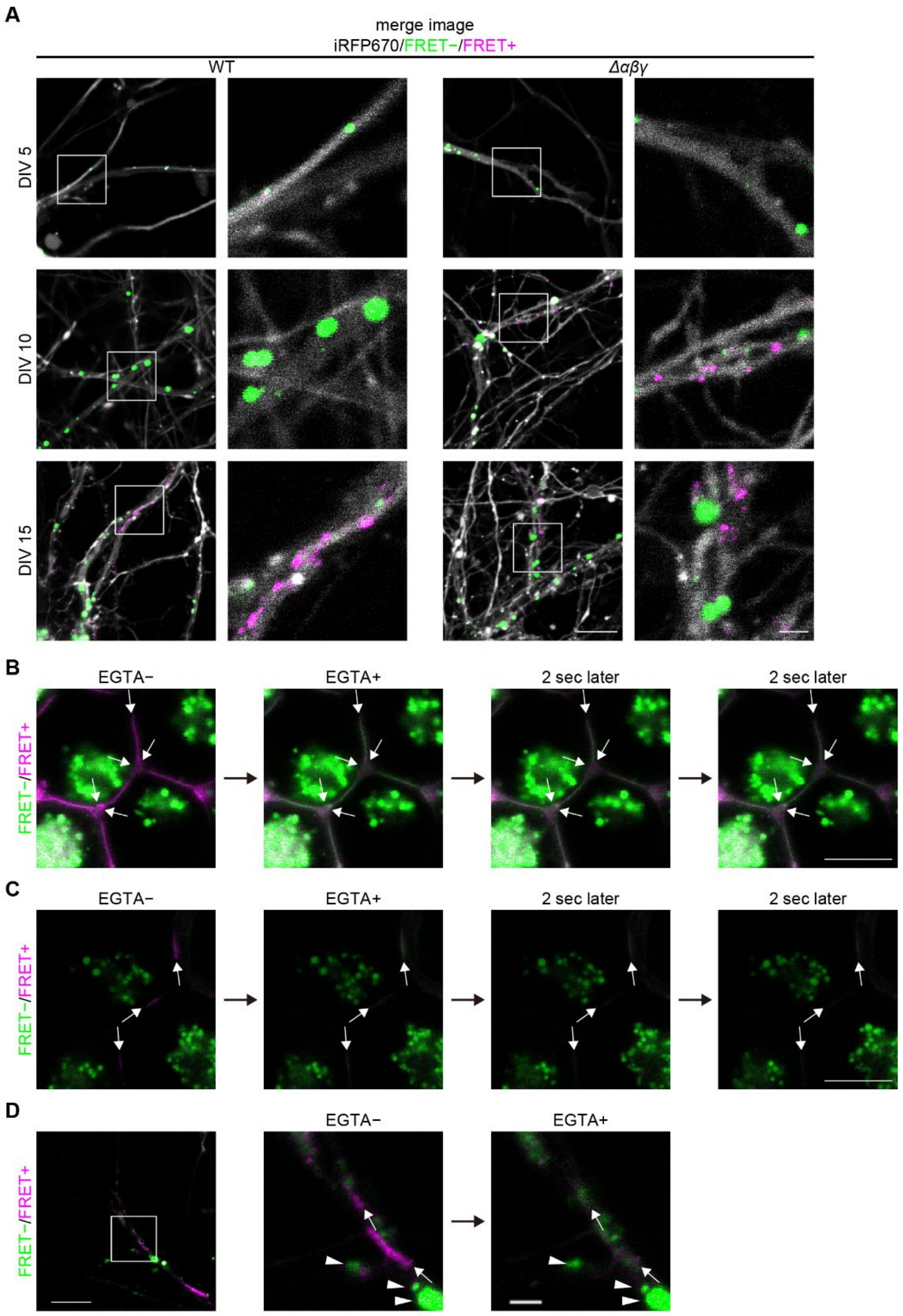


**Fig. S5. Merge and magnified images for Figure 5 B and EGTA-dependent reaction of the γB2 homophilic interaction, related to Figure 5.**

(A) Merge images for Figure 5B. The right panels for each DIV and genotype are magnified images in white squares in the left panels. Scale bar, 10 µm for left, 2 µm for right.

(B) Time-lapse imaging for strong contact sites of combined γB2ΔICD-FRET probe-expressing K562 cells before and after EGTA addition. Several seconds after the addition of EGTA, the combined γB2-FRET probe localizes at contact sites. Scale bar, 10 µm
(C) Time-lapse imaging for weak contact sites of combined γB2ΔICD-FRET probe-expressing K562 cells before and after EGTA addition. Several seconds after the EGTA addition, the combined γB2ΔICD-FRET probe cannot be detected at contact sites. Scale bar, 10 µm
(D) Time-lapse imaging for weak contact sites of combined γB2ΔICD-FRET probe-expressing neurons before and after EGTA addition. The dotted localizing combined γB2ΔICD-FRET probe cannot be detected at contact sites immediately after EGTA addition, but the lined localizing combined γB2ΔICD-FRET probe localizes at contact sites. Scale bar, 10 µm for the left panel, and 2 µm for the right panel.

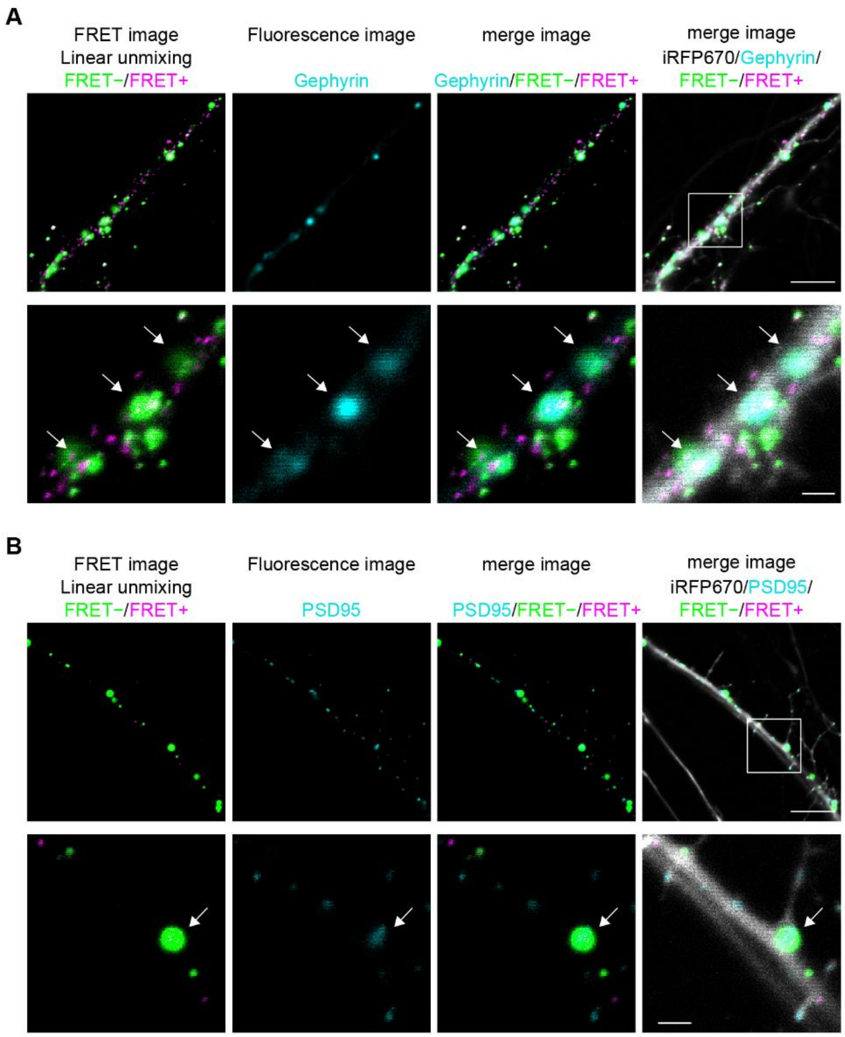


**Fig. S6. Another representative image of the γB2-FRET observation with synaptic markers, related to Figure 6.**

(A) Linear unmixed images, FingR (GPHN)-Cherry, and iRFP670 are shown. The bottom panels are magnified images for the white square of the top panels. Arrows indicate the dots of the combined γB2FL-FRET probe colocalizing with gephyrin. Scale bar, 10 µm for the top and 2 µm for the bottom.

(B) Linear unmixed images, PSD95-TagRFP and iRFP670 are shown. The bottom panels are magnified images for the white square of the top panels. Arrows indicate the dots of the combined γB2FL-FRET probe colocalizing with PSD95. Scale bar, 10 µm for the top and 2 µm for the bottom.

**Table S1. Primers used for qPCR, generation of Pcdhγ floxed mice and generation of PcdhγB2 FRET mice.**

| | |
|---|---|
| B2m-real-F | CCCTGGTCTTTCTGGTGCTT |
| B2m-real-R | TGTTCGGCTTCCCATTCTC |
| Venus-gB2-F | CCCACCCCCTGCTAGCTC |
| FRET-EC5-qPCR-R | CTGCTCGTGGTCGAAGGC |
| gCR1A-F | GTCGACATAACTTCGTATAATGATTATGCTATACGAAGTTATGTTAGTCTCCGGGTTGGTTC |
| gCR1A-R | AAGCTTGTGCCTGGGTGAATCTTGCT |
| gCR1B-F | GGTACCACAGGCCATTGTGAGGCATG |
| gCR1B-R | GTCGACTTGCTGAAGAAACGGATCCC |
| gCR1C-F | GGATCC CTCTTCCTTCTCCCAGCTAC |
| gCR1C-R | GAGCTCACAGGTGTAAGGGATGGAGA |
| gCR1D-F | AAGCTTTAGCCACTAAGCTTTCCTGGG |
| gCR1D-R | GCTAGCAACAAAAGGGTAGCCCCCC |
| gCR1E-F | GCTAGCGCCTTGAGAGCTGACTTCCA |
| gCR1E-R | GCATGCCATGCCCCTTGAATCCAATTC |
| ProbeA-F | TGCGCTTGGGCAAGTTAG |
| ProbeA-R | CAGCCTATAAGAAGCGCTGC |
| ProbeB-F | GACAGACAGAAACAGGGAGC |
| ProbeB-R | CAGCAGTTGCCCAAGCTC |
| pCX1624-F1 | CTAGAGCCTCTGCTAACCATGTTCATGC |
| FRET-R | GTTTAAATCACACCACATCATCTTCGGC |
| F/pBAC-4 | GGGTCGACCTCGAAACTTGTTTATTG |
| R/pBAC-3 | TCCTACAATGTCAAGCTCGACCGATG |
| RP23-244D9 end F1 | CTCTGACCTTCCCTATCACCTTCCTG |
| pBAC-R2 | CGTTTCGATCCTCCCGAATTGACTAGTG |
